# Yap is a Nutrient Sensor Sensitive to the Amino Acid L-Isoleucine and Regulates Expression of Ctgf in Cardiomyocytes

**DOI:** 10.1101/2024.04.25.591002

**Authors:** Victoria L. Nelson, Ashley L. Eadie, Malav Madhu, Mathew Platt, Angella Mercer, Thomas Pulinilkunnil, Petra Kienesberger, Jeremy A. Simpson, Keith R. Brunt

## Abstract

Myocardial infarction and reperfusion is a complex injury consisting of many distinct molecular stress patterns that influence cardiomyocyte survival and adaptation. Cell signalling that is essential to cardiac development also presents potential disease-modifying opportunities to recover and limit myocardial injury or maladaptive remodelling. Here we hypothesized that Yap signalling could be sensitive to one or more molecular stress patterns associated with early acute ischemia. Yap, not Taz, patterns of expression differ in post-myocardial infarct compared to peri-infarct tissue suggesting cell-specificity that would be challenging to resolve for causation in vivo. Using H9c2 ventricular myotubes in vitro as a model, Yap levels were most sensitive to nutrient deprivation compared to other stress patterns typified by ischemia within the first hour of stress. Moreover, this is mediated by amino acid availability, dominantly L-isoleucine, and influences the expression of Ctgf—a major determinant of myocardial adaptation after injury. These findings present novel opportunities for future therapeutic development and risk assessment for myocardial injury and adaptation.

## Introduction

Despite extensive research, myocardial infarction (MI) continues to be a leading cause of morbidity and mortality worldwide[1]; thus, identifying mechanisms of cardiomyocyte adaptation after MI is imperative to the discovery of new therapeutic strategies. MI can be characterized as either reduced or occlusion of blood flow in a coronary artery causing ischemia—the deprivation of oxygen/nutrients and accumulation of metabolic waste causing oxidative stress, cell death and inflammation [2]. Origins of ischemia are diverse and include: coronary vasospasm, the build-up of atherosclerosis coronary plaques, their rupture/erosion, or cardiac microvascular dysfunction and rarefaction (loss or extinction of small blood vessels and their collateralization). Occlusive ischemia of the myocardium can be lethal if blood flow is not rapidly restored [3]. Even with reperfusion, there is an associated injury that can irreversibly compromise cardiac function and then progress to heart failure or arrhythmia morbidity. Current therapeutic interventions (e.g angiotensin II receptor blockers, angiotensin-converting-enzyme inhibitors, beta-blockers or anti-platelet/coagulants) reduce cardiac workload or increase coronary perfusion [2,4] yet therapeutics to deliberately promote adaptive cardiac remodelling post-MI are still required to improve patient outcomes.

Various cells coordinate mechanisms to maintain the contractile capacity and structural integrity of the heart to keep up with the metabolic demands of the body. Many molecular mechanisms are perturbed by cardiac ischemia and resolve on a distinct timeline for each cell type with pathophysiological consequences. Cell survival mechanisms are some of the first adaptations to ischemia, that resist necrosis and/or apoptosis of the cardiomyocytes post-MI [5]. Compensating for the cell loss is critical to preserve mechanical and contractile functions to maintain cardiac output, with fibrotic stabilization and then hypertrophy of the surviving cardiomyocytes being central to short-term adaptation in the infarct and peri-infarct regions of a ventricular free wall under pressure/strain [6]. Cardiac hypertrophy limits wall stress and maintains function by increasing myofilament number and efficiency, without cell proliferation [7]. Initially beneficial, wall chamber shape can elongate with hypertrophy being eccentric and less contractile force—such maladaptive remodelling and ultimately leads to heart failure [6]. What ischemia triggers early in cardiac molecular signalling after MI can therefore reveal potential early activation or repression patterns that might be useful to inform interventions.

Pathways regulating cardiac development likely present opportunity to improve cardiomyocyte adaptation post-MI. Of these pathways, Yap— a key component to the Hippo signalling pathway— serves a direct role in the size regulation of the heart during development [8]. Yap signalling is regulated by several extrinsic and intrinsic stimuli that can initiate the cascade, and generally considered part of mechanical cell signaling, which would be relevant to late remodelling of a dilating ventricle [9]. What early sensitivity Yap signaling has to acute ischemia in unknown. Canonically, when Yap is unphosphorylated it translocates to the nucleus where it binds to a transcription factor(s), predominantly TEAD, to initiate transcription [10]. When Yap is phosphorylated, it is retained in the cytoplasm and, in some cases, this leads to proteasomal degradation [11,12]. Yap signalling is involved in the tissue repair process in other organs (i.e. intestines, liver) and is important in maintaining tissue homeostasis and promoting regeneration after injury [13,14]. There is evidence that active Yap reduced cardiac scar formation and ultimately improved cardiac function [15–17], nevertheless, the cell types and/or underlying triggers from ischemia remain unclear. Unlike, other organs the heart has limited regenerative potential due to the inability to efficiently make new cardiomyocytes [18]. Therefore, exploring the role of cardiac Yap signalling in response to MI and the underlying mechanisms of ischemia affecting Yap could improve existing cardiac remodelling post-MI or provide new possibilities for promoting the adaptive responses of cardiomyocytes.

In the present study, we sought to identify the role of Yap signalling in response to early ischemia stressors and drivers of cardiac injury with cardiomyocyte-like cells modeling responces to early ischemic stress patterns that could influence later remodelling. By modelling the component stressors related to ischemia, we identified that Yap signalling is particularly sensitive to nutrient deprivation, regulated in part by L-isoleucine. Furthermore, the expression of a Yap-associated Ctgf gene was significantly increased consequently to this stimulus in the first hour of nutrient deprivation. These results, establish that Yap signalling alterations are metabolically sensitive and that Yap-regulated signalling in the infarcted myocardium is immediate with associated gene expression changes that could affect longer-term remodelling and adaptation, presenting a novel target for therapy or risk assessment.

## Methods

### In Silico Analyses

The Gene Expression Omnibus (GEO) created by the National Center for Biotechnology Information (NCBI) is a resource for high throughput datasets that follow the NCBI’s guidelines for Minimum Information About a Microarray Experiment (MIAME) [19]. Microarray data sets related to MI were found using the GEO. Gene expression was investigated in the left ventricle of male Wistar rats that received either small, moderate, or large MIs by proximal left coronary artery ligation (Series Accession GDS4907). Gene expression was compared to sham animals (same surgery without ligation). The data presented is relative mRNA (log2-transformed) [20].

### Animal Care

All experimental procedures and animal care were done per Canadian Council on Animal Care guidelines and approved by the University of Guelph’s Animal Care Committee. Animals were provided with food and water *ad libitum* and maintained on a light:dark (12:12 hour) cycle.

Volume Overload Heart Failure (AMI)

### Pre-clinical Models

Acute myocardial infarction was attained by permanent ligation of the left anterior descending (LAD) coronary artery in C57BL/6 adult male mice (8 weeks of age). Animals were anesthetized (2%:100% isofluorane:O 2), intubated and ventilated using the Harvard Apparatus at 200 breaths/minute. Para-sternal thoracotomy was done under sterile surgical conditions, next a 7-0 Surgipro**^TM^** II polypropylene suture (Covidien) was used to ligate the LAD below the atrioventricular border; confirmed with blanching of the wall below the ligation. The same operation was performed for Sham animals, but no ligation was made. After surgery, animals were checked twice daily for any complications. AMI animals and shams were humanely harvested at 7-& 28 days post-injury to assess expression in remodelling phase post-MI.

### Tissue Collection and Preparation

Hearts were excised and immediately flash frozen and stored at -80°C until use. Samples were ground using mortar and pestle while in liquid nitrogen. 10-15mg of the sample was homogenized for 30 seconds in 120μl NP-40-based lysis buffer (1% NP-40), then placed on ice for 30 minutes. Fresh lysis buffer was made by adding a 1:100 phosphatase inhibitor (PHI) cocktail (524628, EMD Millipore), a protease inhibitor (PI) (m250, VWR International) and activated sodium orthovanadate (Calbiochem). Samples were centrifuged at 4°C for 2 minutes at 2000rcf, and the supernatant was transferred to a new 1.5mL tube, and centrifuged again at 4°C for 30 minutes (1200 rcf). Using a 28 ½-guage insulin syringe (Becton Dickinson) the supernatant was transferred to a fresh 1.5mL tube.

### Immunohistochemistry

Hearts were fixed in diastole using 1xPBS, 50mmol KCL perfusion followed by immersion in 10% neutral buffered formalin (VWR International) overnight. After being placed in a tissue cassette were dehydrated in 70,80, and 100% ethanol sequentially, cleared in Xylene, and infiltrated with paraffin wax using an automated tissue processor. Samples were blocked in paraffin wax until hardened and positioned to allow cross-sectional cuts at 4um on a microtome. Paraffin sections were deparaffinized and rehydrated using xylene, along with ethanol. The sections were rinsed in tap water for 5 minutes followed by incubation in Tris-EDTA (pH 9) for 15 minutes using a steamer. They were then incubated in BLOXALL (Vector Labs) Blocking Solution for 10 minutes. Sections were washed in 1XPBS for 5 minutes and incubated for 20 minutes with prediluted normal blocking serum (Vector Labs). The sections were incubated for 30 minutes with 60μl Yap/Taz primary antibody (8418, Cell Signaling Technologies); diluted (1:200) in a PBS-T/ 20 bovine serum albumin (BSA) (1X PBS, 0.1% Tween 20, and 1% BSA solution). Sections were then washed in PBS for 5 minutes. Prediluted biotinylated secondary antibodies (Horse Anti-Mouse/Rabbit IgG; VectorBioLabs) were incubated for 30 minutes. The sections were washed in PBS for 5 minutes and incubated for 30 minutes in R.T.U.

VECTASTAIN Elite ABC Reagent (Vector Labs,). The sections were then washed for 5 minutes in PBS and stained with DAB Peroxidase Substrate Solution (Vector Labs), for 3 minutes under the microscope. The sections were quickly rinsed in tap water, counterstained in Hematoxylin (Vector) for 1 minute, and washed in tap water for 5 minutes. Sections were dehydrated in reverse sequential ethanol and xylene solutions and then mounted using DPX Mountant for Histology (Sigma-Aldrich) and allowed to dry overnight.

### Cell Culture

H9c2 embryonic rat cardiomyoblasts from American Type Culture Collection, were expanded from a cryostock, in Dulbeco’s Modified Eagle Medium (DMEM; Gibco®) supplemented with 10% Fetal Bovine Serum (12483020; FBS, Gibco), in 37°C, 95% air, and 5% CO^2^ incubation. Cells were cultured to 80% confluency before being passaged. The passage number did not surpass 18. For experiments, cells were seeded at 500,000 cells/60mm plate.

### Cell Stress Treatments

Differentiation: H9c2 cells were differentiated to quiescent myotubes with the removal of FBS for 6 days, with media changes every 72 hours (DMEM, no FBS). Cells were treated for 1hour acutely under the following conditions:

Nutrient Deprivation Solution: A 10nM Hepes (Ambresco) was made in Earle’s Balanced Salt Solution (EBSS, Gibco) and cells were treated for one hour. The solution was kept at 4°C for up to a year.

Cytokine Stress: Tumor necrosis factor-alpha was supplemented (20mg/mL) into DMEM. Mitophagic Stress: Carbonyl cyanide-p-trifluoromethoxyphenylhydrazone (FCCP) [2μM] was prepared in DMEM from a 100mM DMSO stock solution.

Hypoxia: Cells were incubated in DMEM at 1% O^2^ using a HERAcell 150.i CO^2^ incubator (Thermo Scientific).

Oxidative Stress: Hydrogen peroxide was diluted from a 0.9M stabilized stock solution, diluted in PBS to 0.1M, then prepared in DMEM and cells were treated for 24 hours at [450μM].

Ischemia: Cells were incubated at 1% O^2^ in DMEM without glucose for 24 hours.

L-isoleucine: L-isoleucine (J63045.14, Thermo Scientific) powder was added directly to our nutrient deprivation solution to a final concentration of 105.0 mg/L, vortexed and stored at 4°C.

L-valine: L-valine (J62943.06, Thermo Scientific) powder was added directly to our nutrient deprivation solution to a final concentration of 105.0 mg/L, vortexed and stored at 4°C.

L-Threonine: L-threonine (J63709.30, Thermo Scientific) powder was added directly to our nutrient deprivation solution to a final concentration of 105.0 mg/L, vortexed and stored at 4°C.

### Resazurin Metabolic Assay

Following treatment, 10μl 10% PrestoBlue Cell Viability Reagent (A13261, Thermo Scientific), was added to each well of the plate and was incubated at 37°C for 1 hour. Fluorescence was measured with excitation (560nM) and emission (590nM) wavelengths. The 560/590nM control well (no cells) was subtracted from its respective treatment group when calculating metabolic activity.

## Immunoblotting

### Cell Collection and Whole Cell Lysate Preparation

Media was aspirated, cells rinsed with PBS and scraped in an NP-40-based lysis buffer containing 1:100 of PI, PHI, and sodium orthovanadate stored at -80°C in 1.5mL tubes. The lysates were sonicated for 10 seconds at 20kHz and 30% amplitude. Protein concentration was quantified using a bicinchoninic acid assay kit.

### Nuclear and Cytoplasmic Protein Extraction

After treatment media was aspirated from the cells, rinsed with PBS and then a cell scraper was used to collect into 60μL of hypotonic buffer (10mM Hepes,10mM KCl,0.1mM EDTA, 0.1mM EGTA,1mM DTT at pH7.9) supplemented with a 1:100 ratio of PHI, PI, and sodium orthovanadate. Lysates were on ice for 20 minutes, followed by centrifugation at 20,000xg for 5 minutes. The supernatant was collected as the cytosolic fraction. The pellet was resuspended in 120μL hypotonic buffer and centrifuged again; supernatant was collected as wash. The pellet was resuspended in 30μL hypertonic buffer (20mM Hepes, 0.4M NaCl, 1mM EDTA, 1mM EGTA, 1mM DTT) and left on ice for 30 minutes, followed by centrifugation for 5 minutes at 10,00xg. Supernatant was collected as nuclear fraction. Protein concentration was quantified using a Bradford Assay kit.

### Western Blotting

Lysates were boiled (99°C for 5 minutes) in Laemli buffer with DTT samples loaded beside a Plus Protein Standards Kaleidoscope™ ladder (BioRad) into a 3.5% 1X stacking gel. Samples were resolved in a 10% Mini-Protean Gel at 90V in 1X Tris/Glycine/SDS Electrophoresis Buffer (1610772; BioRad) until the dye front reached the bottom of the gel. Then, samples were transferred onto a nitrocellulose membrane (0.2μM, 1620112; BioRad) at 100V for 75 minutes at 4°C in a 1X Tris/Glycine Transfer Buffer (1610771EDU; BioRad). After transfer, the membrane was washed with ddH^2^O then stained and imaged using Reversible Protein Stain (24580, Thermo Scientific) on the ChemiDoc MP Imaging System (BioRad). The stain was removed using Memcode Eraser (24580, Thermo Scientific) and rinsed in ddH^2^O followed by 1X Tris-Buffered-Saline-Tween 20 (TBST). Membrane was blocked for 1 hour in 5% skim milk/1X TBST and rinsed in TBST before being incubated overnight at 4°C in primary antibody. p-Yap S127 (1:1000, #13008, Cell Signaling), p-Yap S397 (1:1000, 13619, Cell Signaling) Yap/Taz (1:1000, 8418, Cell Signaling) Lamin A/C (1:1000, 2032, Cell Signaling Technologies), GAPDH (1:1000, TA802519, Origene). After membranes were washed in TBST and incubated in horseradish peroxidase-conjugated secondary antibody anti-rabbit (1:1000-1:2000, 31460, Invitrogen) or anti-mouse (1:1000-1:2000, Biorad) with 5% milk for 2 hours at room temperature. Clarity Western ECL Substrate (1705060S; BioRad) luminol/enhancer and peroxide solutions were used to image membranes. Between targets, membranes were stripped in 25mL of 0.5M Tris-HCl/SDS buffer with 125μl β-mercaptoethanol for 45 minutes followed by blocking. Densitometric calculations were done using ImageLab Software v5.0 (BioRad), and all targets were normalized to total protein density from the respective Memcode lane.

### Isolation and Measurement of Free Amino Acids

1 million cells were suspended in 60 μL of MilliQ water and 60μL of 2M perchloric acid (CA71007-908, VWR) along with 120μL of internal standard (4 μg/mL) containing arginine-d7, glycine-d5, lysine-d4, and leucine-d3 (made from: CDN Isotopes, D-7786, D-0277, D-2554, and D-1973 respectively) followed by being vortexed and lysed using an ultra sonicator for 10 seconds. Followed by two minutes of sonication at room temperature followed by 5 minutes in an ice bath twice to precipitate protein, then centrifuged at 4°C for 15 minutes at 13,000rpm. The supernatant was transferred to a clean tube and the cell pellet was washed with 60μL MilliQ water. 150μL of supernatant was neutralized with 120-150μL of potassium hydroxide [2M] (CABH9262, VWR) and then centrifuged at 13,000 rpm. Supernatant was transferred to a new tube and the extract was freeze-dried for two to four hours, before being resuspended in 60μL of 50:50 Methanol: MilliQ water. Resuspended extract (10μL) was transferred to an autosampler vial and mixed with 70μL of Borate Buffer (186003836, Waters) for 5 minutes as per the Waters AccQTag Derivatization Kit. Samples were vortexed then 20μL AccQTag Derivatization Agent (186003836, Waters) was added followed by another vortex and left to sit for one minute before being placed in the heating block for 10 minutes at 55°C then vortexed. Samples were run using the Waters Acquity ultra-performance liquid chromatography (UPLC), Xevo-TQS-micro–Tandem Mass Spectrometer using Multiple Reaction monitoring for each BCAA with respective internal standards. Results were quantified using TargetLynx (Waters) software, samples were corrected by milligrams of protein.

### Quantitative Polymerase Chain Reaction (qPCR)

Cell harvesting and RNA isolation were done using the Qiagen RNeasy Mini kit (74106, Qiagen) and then stored at -80°C until use. A 20μl cDNA reaction mixture was prepared by adding sample-specific volume equivalency of 4μg RNA and topped up to the 20μl volume with Nuclease-free H^2^O into RNase/DNase-free strip-tubes along with 20μl 2X Master Mix (43-688-14, Fisher Scientific). The strip tubes were briefly centrifuged then placed in the Mastercycler Nexus Gradient Thermocycler (Eppendorf) for 10 minutes at 25°C, 37°C for 120 minutes, 85°C for 5 minutes, with a 4°C hold. Samples were stored at -80°C until use.

Primers were designed in OLIGO Primer Analysis Software V6.31 (Molecular Biology Insights, Inc.). Each primer set was made into a qPCR master mix, containing 5μl Platinum SYBR Green qPCR Supermix-UDG (11733038, Invitrogen), 0.2μl forward primer (10mM), 0.2μl reverse primer (10mM), 0.02μl ROX reference dye, and 2.58μl Nuclease-free H2O. After 8μl of the Master Mix was added to each well in a 96-well qPCR plate, along with 2μl of cDNA. No-template control, containing only the master mix was run at the same time. The plate was sealed, then centrifuged for 3 minutes at 2500rpm. The reaction was incubated at 50°C for 120s, 95°C for 120s, and 35-40 cycles of incubation at 95°C for 3s and 60°C for 30s (Applied Biosystems Viia 7). All qPCR data was analyzed as per Minimum Information for Publication of Quantitative Real-time PCR Experiments (MIQE). Two reference genes were used that were not influenced by culture conditions or treatments for group effect comparison (Supplemental Fig.3). Data was analyzed using Igfbp3 as a reference gene and confirmed by a second reference gene, Hprt1 to compare nutrient deprivation to DMEM controls and Igfbp3 and B2m as reference genes for comparison between nutrient deprivation + L-isoleucine and DMEM controls.

### Immunocytochemistry

Cells were seeded (600,000/60mm plate) on a glass coverslip and differentiated (protocol above), followed by respective treatments (control vs. nutrient deprivation). Cells were washed with 1XPBS and then fixed using warmed formaldehyde (4% in PBS) for 5 minutes. The cells were washed three times with ice-cold PBS before being permeabilized in 0.1% Triton-X-100 for 15 mins then washed again three times in PBS. Cells were then blocked using a 1% BSA in 1XPBS 0.1% Tween 20 solution for 15 minutes, followed by another wash. Cells were then incubated in AlexaFluor 488 Phalloidin (Thermo Scientific, diluted 1:1000) stain in 1XPBS for 30 minutes while protected from light; then washed and incubated in primary antibodies diluted 1:400 at 4°C for 24 hours. After a wash they were incubated in a fluorophore-conjugated secondary antibody (Alexa Fluor 647 donkey-anti-rabbit, diluted 1:400) for 30 minutes, followed by a wash. Cells were then stained with Hoechst 33342 which was diluted 1:3000 in 1XPBS for 1 minute, and washed one last time before the coverslip was mounted to a glass slide using 1:1 glycerol and PBS, then sealed with clear nail polish (at the edges) and allowed to dry overnight at 4°C.

### Statistical Analysis

All results are representative of the mean ± standard deviation. Statistical analysis was done using GraphPad Prism 9 (GraphPad Software Inc.). Pairwise comparisons were done using a student’s two-tail t-Test, while any results that had three or more groups were analyzed using a one-way or analysis of variance (ANOVA). Statistical significances were measured using Tukey’s post-hoc test. To be considered statistically significant p-values <0.05.

## Results

### Yap Signalling in Myocardial Infarction and Associated Stressors

To establish the effects of MI on Yap signalling in the cardiac remodeling, immunohistochemistry was performed on 28-day post-MI mouse hearts and compared to shams. Positive Yap-staining (brown/red colouring) was observed and localized mainly in the peri-infarct zone post-MI (Fig. 1 I-A/B) compared to the remote region and to some extent in the remote ventricles (Fig1 I-C/D). An increase in Yap levels was detectable by western blot as soon as 1-week post-MI in the peri-infarct zone with no change in the infarct zone, suggesting it is regulating early and late remodelling. Taz levels remained unchanged in the peri-infarct and infarct zones compared to shams (Fig1 II). To determine how Yap changes in response to specific MI stress patterns associated with ischemia, we used cardiomyocyte-like (H9c2) cells as a model and screened several commonly associated drivers (nutrient deprivation, cytokine, mitophagic, hypoxic, oxidative, and ischemic stress). We found that total Yap expression was increased in differentiated cells compared to proliferative cells, however, Yap levels were unchanged between differentiated cells and most other ischemic stressors/stimulus conditions (Fig. 1-III). All conditions were compared and tested using differentiated cells as they are the most predominant state of cardiomyocytes in a developed heart. As phosphorylation is important for Yap signalling, we examined one of the key phosphorylation sites (p-S397) in our various conditions. We determined that of all stimuli conditions Yap (p-S397) was most significantly increased under nutrient deprivation (acute) conditions compared to differentiated or proliferative cells alone (Fig. 1-IV). Unlike the ischemic stress, which is low nutrient, this total nutrient deprivation models a severe condition expected under total occlusion. The increase in Yap (p-S397) was confirmed by the ratio of Yap (p-S397) to total Yap (Fig. 1-V). These findings suggest that Yap signalling is most sensitive to severe metabolic stress associated with MI and that nutrient deprivation has a large impact on the phosphorylation of Yap. Many phosphorylation sites exist in the Yap signalling pathway that are important for determining whether Yap will translocate to the nucleus (de-phosphorylated) or be retained in the cytoplasmic and/or degraded (phosphorylated) of these is Yap (p-S127) which was also significantly increased compared to differentiated controls (Supplemental Fig.1)

**Figure 1:**
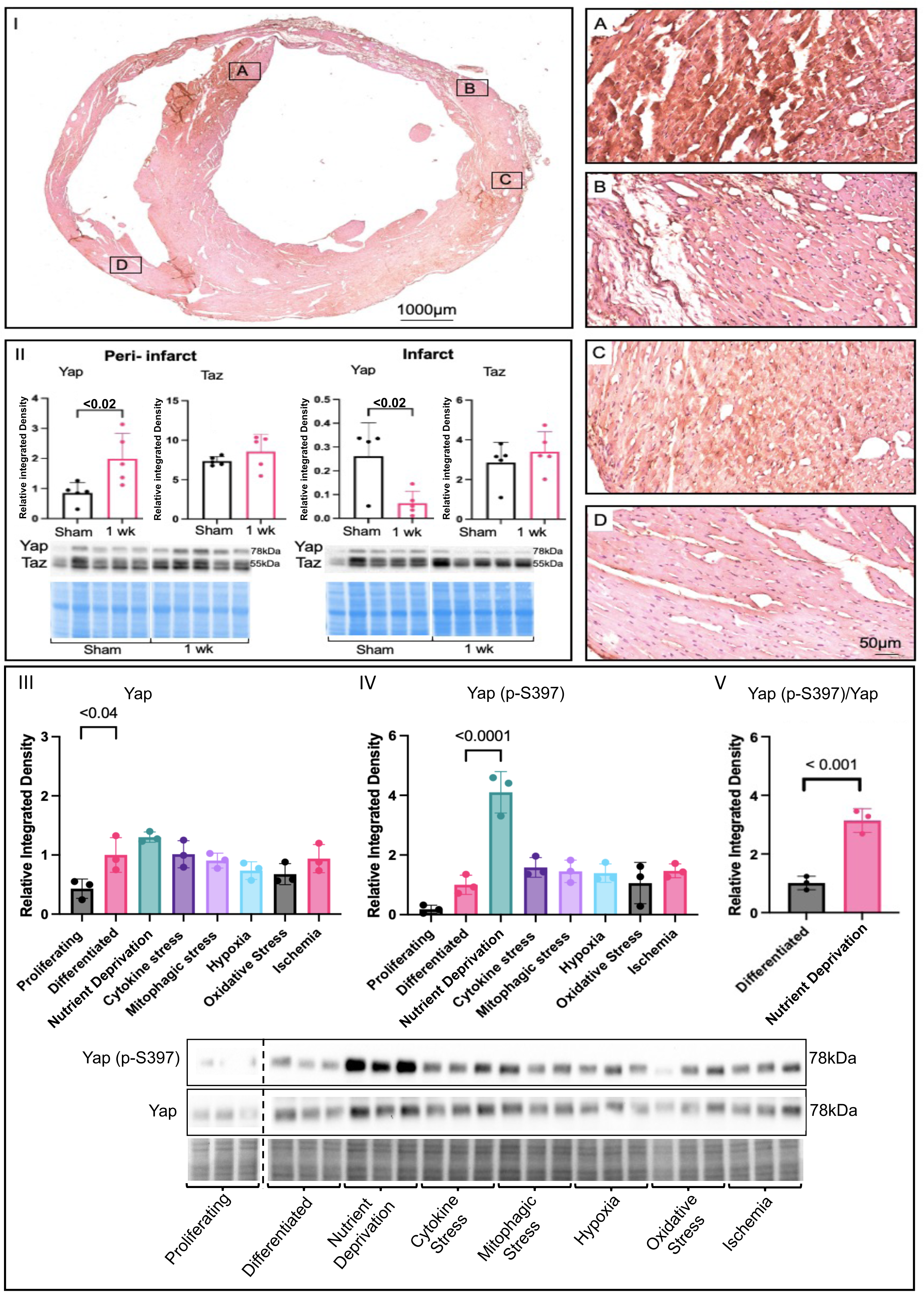
Changes in Yap Signalling Under Ischemic Conditions. **(I)** Yap immunohistochemistry of a 28-day post MI mouse heart. Peri-infarct zone **(A-C)** had higher Yap expression seen by the brown positive Yap signal, compared to the remote region **(D)** mostly pink and controls (not shown). **(II)** Yap expression was increased in the peri-infarct zone and decreased in the infarct zone of 1 week post MI mice compared to shams. Immunoblot and densitometric analysis of Yap and Taz peri-infarct and infarct zone of sham compared to 1 week post MI mice. **(III)** Analysis of Yap in screen of proliferating (DMEM+10% FBS), differentiated (DMEM), metabolic shock (EBSS, 1h), cytokine stress (TNF⍶ 20ng/mL, 12h), mitophagic stress (FCCP 2µM, 12h), hypoxia (O_2_ 1%, 24h), oxidative stress (H_2_O_2_ 450µM, 24h), ischemia (O_2_ 1%, + 0% glucose, 24h) in H9c2 cells where expression was increased in differentiated cells compared to proliferating. **(IV)** Yap (p-S397) was significantly increased under metabolic shock conditions compared to all other ischemic stressors. **(V)** Significant increase in ratioed Yap (p-S397) to total Yap in nutrient deprivation conditions with corresponding relative blots and Pierce® Memcode to demonstrate uniform loading.

### Nutrient Deprivation Effects Yap Post-Translation Modification & Compartmentation

To investigate whether nutrient deprivation affects the translocation of Yap or phosphorylated Yap we used immunohistochemistry with phalloidin which stains f-actin on the cellular membrane (green), Hoechst for nuclei staining (blue) and Yap fluorescence antibody (red). Total Yap levels did not change and there was little indication of migration or translocation to the nucleus compared to controls. Yet, both Yap (p-S397) and Yap (p-S127) appear to cluster towards the nucleus (speckling and purplish colour of nuclei) in nutrient deprivation conditions compared to controls (Fig. 2-I). To substantiate the translocation of Yap and phosphorylated Yap to the nucleus, nuclear and cytoplasmic fractionation were performed. There were no changes in total Yap levels in either fraction compared to controls which agrees with immunohistochemistry results. However, Yap (p-S397) was significantly increased in the nucleus and the cytoplasm, while Yap (p-S127) was increased only in the nucleus compared to controls (Fig. 2-II) suggesting that acute nutrient deprivation plays a role in the translocation of phosphorylated Yap.

**Figure 2.**
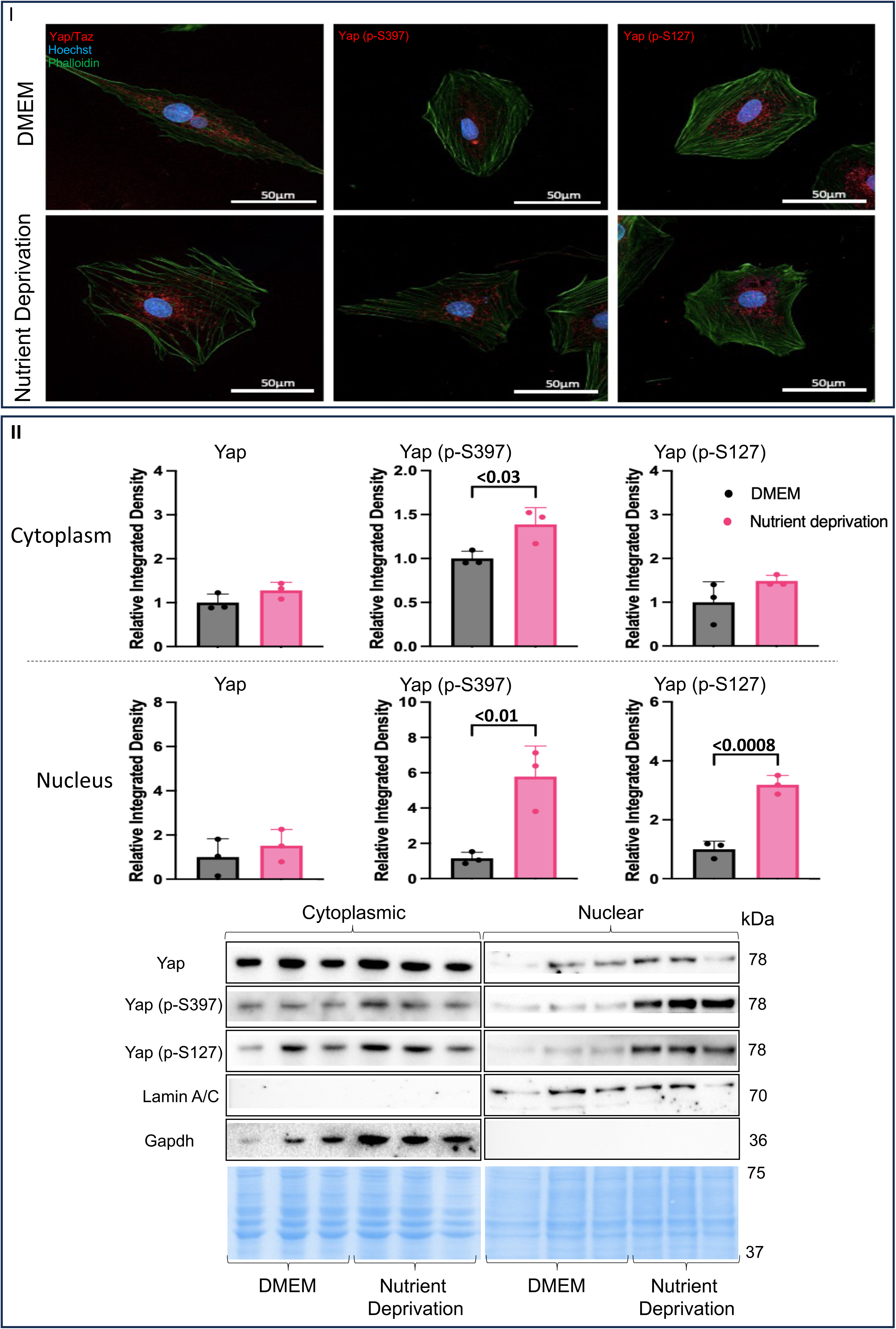
Evidence of Nuclear Compartmentalization Under Nutrient Deprivation Conditions. H9c2 cells were differentiated for 6 days in the absence of FBS, on the day of the experiment controls received a media change and our treatment groups received nutrient deprivation solution for 1 hour. Cells were permeabilized and blocked followed by staining with Phalloidin, and Hoechst and incubated with respective fluorescent antibodies for immunofluorescent. **(I)** No changes in Yap compartmentalization (nuclear and cytoplasmic) compared to controls. Yap (p-S397) and Yap (p-S127) were increased in the nucleus of metabolic shock conditions compared to controls. **II)** Cytoplasmic Yap expression was unchanged in nutrient deprivation conditions compared to controls. **II)** Yap (p-S397) expression was increased in the cytoplasm, while Yap (p-S127) was unchanged under nutrient deprivation conditions compared to controls. Nuclear Yap expression was unchanged compared to controls. Nuclear Yap (p-S397) and Yap (p-S127) expression were both increased in nutrient deprivation conditions compared to controls. Representative blots for Yap, Yap (p-S397) and Yap (p-S127) and nuclear and cytoplasmic markers, Lamin A/C and Gapdh, respectively and Pierce® Memcode demonstrating uniform protein loading.

### Amino Acids Nutrient Sensitivity Impacts Yap Signalling

Nutrient deprivation is one component of ischemia, along with other components such as hypoxia and oxidative stress (Supplemental Fig. 2-I). Hypoxia and oxidative stress conditions resulted had no effects on Yap signalling, unlike nutrient deprivation (Fig. 1-III, IV) which increased phosphorylated Yap compared to controls. Nutrient deprivation is defined as an acute state where cells are deprived of the three essential macronutrients: glucose, fatty acids, and amino acids. To establish which of the three essential macronutrients was responsible for the increase in phosphorylated Yap we used a reductive reasoning (Supplemental Fig. II-VI) to determine that amino acids were likely driving the changes in Yap signalling under nutrient deprivation conditions. To better understand the effects of nutrient deprivation conditions on amino acids, we characterized the changes in free amino acids in nutrient deprivation conditions compared to controls by UPLC. Free essential amino acid levels were decreased in nutrient deprivation conditions compared to controls, while non-essential free amino acid levels were increased in nutrient deprivation conditions compared to controls (Fig. 3-I). Furthermore, nutrient deprivation conditions decreased all branched-chain amino acids to the same extent compared to controls. To determine whether which amino acids were able to attenuate the increase in phosphorylated Yap in nutrient deprivation conditions compared to controls we chose the three amino acids that were most significantly decreased and supplemented them into the nutrient deprivation conditions. Concentrations equivalent to DMEM control media (0.8mM) of the three amino acids, L-threonine, L-valine, and L-isoleucine were added separately into the nutrient deprivation conditions. Supplementation with L-threonine and L-isoleucine increased total Yap levels, while total Yap levels remained unchanged with L-valine supplementation compared to DMEM controls (dashed line at 1). Furthermore, phosphorylated Yap S397 and S127 were increased with L-valine supplementation compared to controls, while supplementation with L-threonine and L-isoleucine did not change phosphorylated Yap S397 and S127 levels compared to DMEM controls (Fig. 3-II). An attenuation in phosphorylated Yap S397 and S127 with supplementation of L-threonine and L-isoleucine suggests that the absence of these amino acids could be responsible for the increase in phosphorylated Yap levels in nutrient deprivation conditions compared to DMEM controls (Fig. 1-IV). To further narrow down our focus we considered that threonine is catabolized to isoleucine and is regulated by an inhibitory feedback mechanism, where isoleucine binds allosterically to threonine deaminase (the first enzyme in the catabolism pathway) when isoleucine levels are high, therefore inhibiting the binding of threonine. In contrast, when isoleucine levels are low, threonine can bind and the catabolism to isoleucine can continue suggesting that isoleucine is a limiting factor [21]. Since the relationship between threonine and isoleucine relies on isoleucine as a limiting factor, all experiments moving forward were performed with L-isoleucine.

**Figure 3.**
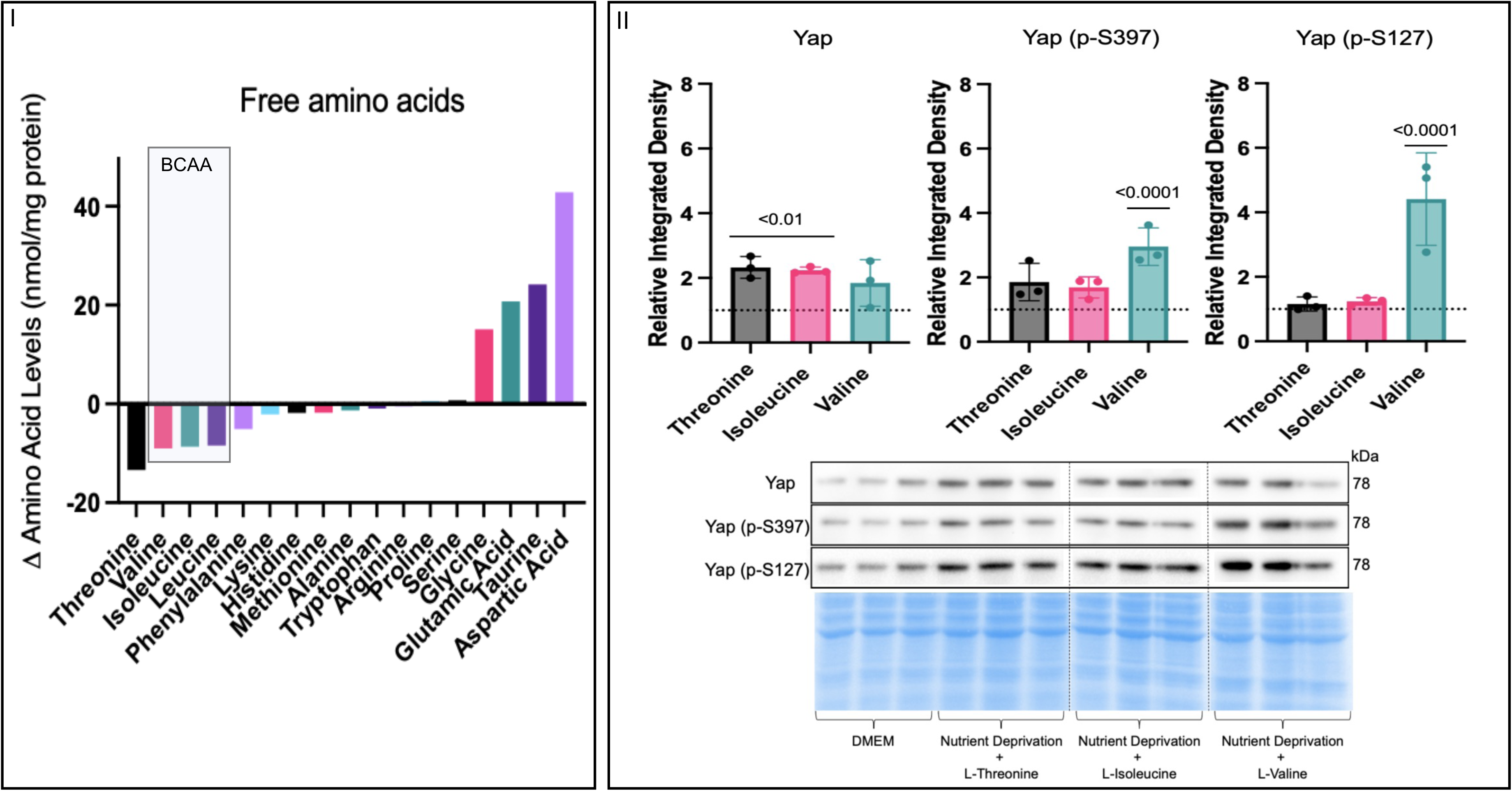
Effects on Free Amino Acids Under Nutrient Deprivation Conditions. H9c2 cells were differentiated for 6 days in the absence of FBS, on the day of the experiment controls received a media change and our treatment groups received nutrient deprivation solution for 1 hour. **(I)** UPLC free amino acid levels differences between controls and nutrient deprivation conditions. All non-essential amino acid levels were increased where all essential amino acid levels were decreased in nutrient deprivation compared to controls. All three branched-chain amino acids were decreased to the same extent. **II)** On the day of the experiment controls received a media change and our treatment groups received nutrient deprivation solution with respective amino acid supplemented for 1 hour before harvest. Total Yap expression was increased in threonine and isoleucine groups compared to DMEM controls (dashed line) while there was no change between controls and valine supplementation. Supplementation of threonine and isoleucine caused no changes in Yap (p-S397) expression compared to DMEM controls. Yap (p-S397) expression was increased in the Valine group compared to controls. Yap (p-S127) expression was unchanged between threonine and isoleucine groups and controls and increased in the valine treatment compared to controls. Representative blots for Yap, Yap (p-S397) and Yap (p-S127) and Pierce® Memcode demonstrating uniform protein loading.

To determine the effects of L-isoleucine supplementation on the translocation patterns of Yap and phosphorylated Yap to/from the nucleus, sub-cellular fractionation was done to isolate the nuclear and cytoplasmic fractions. Interestingly, with additional data points, total Yap levels were increased in nutrient deprivation conditions compared to DMEM controls in the cytoplasmic fraction, though the effect size is small (Fig.4-I). Total Yap levels were unchanged between DMEM controls and the nutrient deprivation + L-isoleucine in the cytoplasm (Fig.4-Supplementation with L-isoleucine attenuated the increase in cytoplasmic phosphorylated Yap S397 levels seen in the nutrient deprivation conditions alone compared to DMEM controls (Fig.4-II). However, supplementation with L-isoleucine did not affect phosphorylated Yap S397 and S127 levels in the nuclear fraction or Yap S127 in the cytoplasm compared to nutrient deprivation alone (Fig. 4-III, V, VI). These findings suggest that L-isoleucine has a major role in the phosphorylation patterns of Yap S397 but does not reverse all effects of nutrient deprivation and its elicited compartmentation.

**Figure 4.**
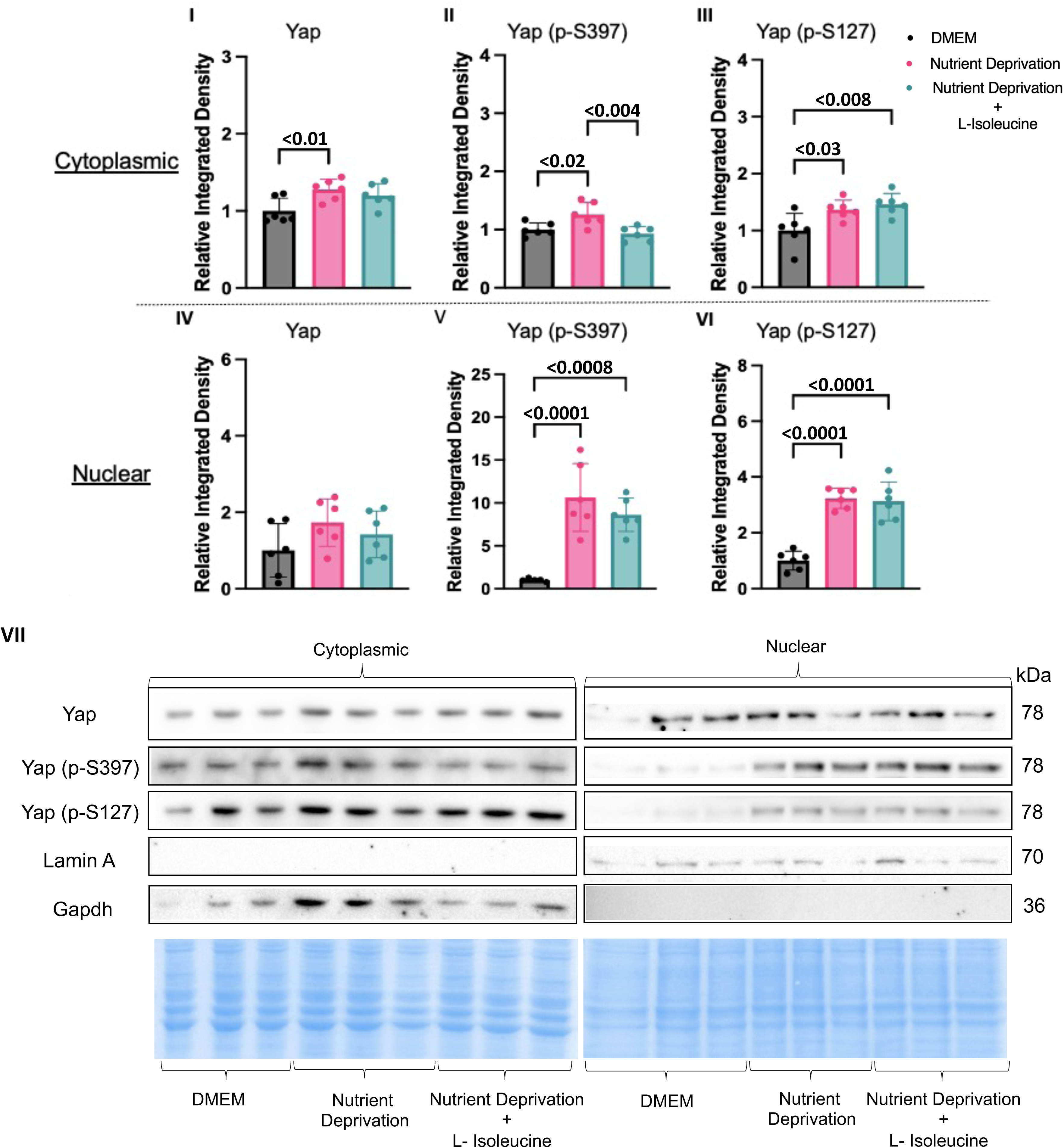
L-Isoleucine Decreased Cytoplasmic Yap (p-S397) Back to DMEM Control Levels. H9c2 cells were differentiated for 6 days in the absence of FBS, on the day of the experiment controls received a media change and our treatment groups received nutrient deprivation solution or nutrient deprivation with L-isoleucine for 1 hour, subsequently followed by nuclear and cytoplasmic fractionation. **(I)** Nutrient deprivation increased Yap expression compared to controls, however, it was unaffected between control and nutrient deprivation with L-isoleucine. **(II)** L-isoleucine supplementation decreased cytoplasmic Yap (p-S397) levels compared to nutrient deprivation alone, while it did not affect cytoplasmic Yap (p-S127) **(III)**, or any of the nuclear fraction for Yap, Yap (p-S397) and Yap (p-S127) **(IV-VI).** Representative blots for Yap, Yap (p-S397) and Yap (p-S127) and nuclear and cytoplasmic markers, Lamin A/C and Gapdh, respectively and Pierce® memcode demonstrating uniform protein loading **(VII)**.

### Yap-associated Gene Target Expression Relevant to MI

The changes in phosphorylation patterns and Yap at the protein level could lead to alterations in downstream Yap-mediated gene expression. Therefore, to investigate the effects of nutrient deprivation and nutrient deprivation + L-isoleucine on gene expression. To simplify a targeted search for qPCR plausible Yap-mediated gene targets we first considered five classical categories of MI recovery: neovascularization, inflammation, cell survival, cardiac performance, and matrix remodelling. We accessed an open-access microarray data set from the Entrez Gene Expression Omnibus (GEO) to screen for any Yap-associated genes in MI, but found only two were listed in the data set (data set: GDS4907), connective tissue growth factor (Ctgf) and c-c motif chemokine ligand 24 (Ccl24). Ctgf expression was significantly increased in the large MI group compared to all other groups, while there was no change in Ccl24 expression between any of the groups (Fig. 5-II). For each MI category, we then performed a simple qPCR screen of two Yap-associated genes that aligned with a category of MI remodelling. Our qPCR results showed no significant changes in any Yap-associated genes related to four of the MI recovery themes, neovascularization, inflammation, cell survival, or cardiac performance. However, as with the GEO data set, Ctgf, a matrix-associated gene, was significantly increased in nutrient deprivation compared to controls. Yet, addition of L-isoleucine did not attenuate Ctgf expression, rather it further increased expression. Collectively, these results suggest that Yap-associated Ctgf expression could be a downstream target in response to nutrient deprivation conditions caused by MI, and proportional to the size/severity of infarction.

**Figure 5.**
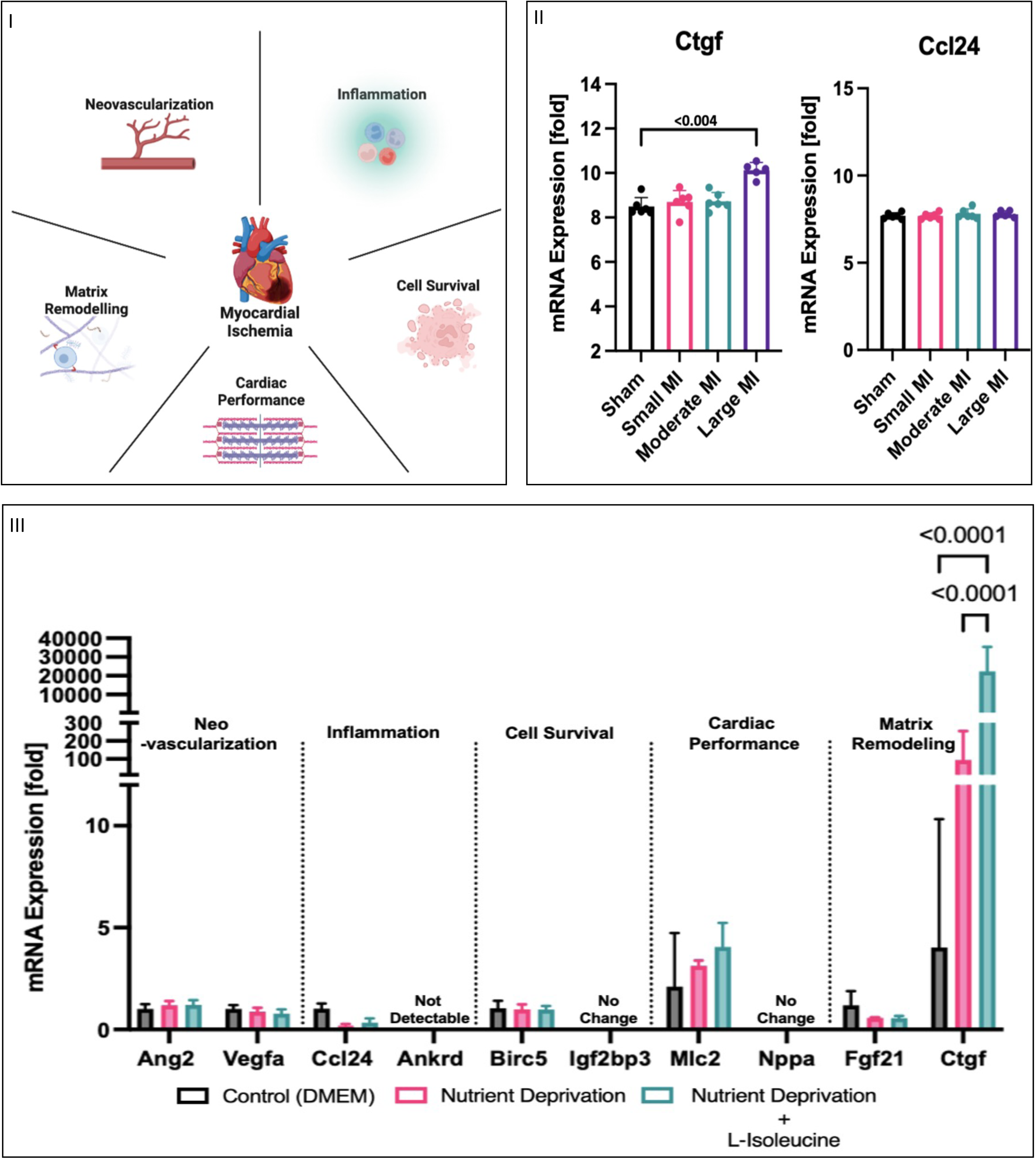
Increase in mRNA Expression of Matrix-Related Genes. I) Myocardial infarction has five categories associated with recovery after a MI: neo-vascularization, inflammation, cell survival, cardiac performance, and matrix remodelling. II) mRNA (log2-transformed) was analyzed in rat MI model using public microarray datasets where, left ventricles were removed from rats who underwent left coronary artery ligation to induce large, moderate, or small myocardial infarctions. Genes associated with Yap were identified as Ctgf, and Ccl24. Ctgf was significantly increased in the large MI group compared to all other groups, while Ccl24 did not vary between all four group isncluding control shams. II**I)** H9c2 cells were differentiated to determine any similar association to AMI after nutrient deprivation with or without L-isoleucine for 1 hour. Two genes that had a priori links to Yap regulation from each MI recovery category were tested, with no changes in genes associated with neo-vascularization, inflammation, cell survival or cardiac performance. However, Ctgf, which is associated with matrix remodelling was significantly increased in nutrient deprivation with L-isoleucine compared to nutrient deprivation alone.

## Discussion

Several studies suggest Yap signalling can improve outcomes after MI [17,22,23], however, the molecular drivers and mechanisms that regulate Yap and its effects in cardiac-related cells post-MI are less understood. The purpose of this study was to determine the stressors and molecular mechanisms that may regulate and/or promote Yap signalling after MI. We observed increased Yap in the infarct and peri-infarct zone after MI, which could be explained by an increase in Yap in cardiomyocytes or fibroblasts after MI, as Yap expression has been seen to increase in cardiac fibroblasts three to five days after MI [24]. Additionally, we found that amongst simplified ischemic stress patterns associated with MI in H9c2 myotubules Yap signalling was particularly sensitive to the deprivation of essential nutrients (glucose, fatty acids, and amino acids). Our data then resolved that this was predominately influenced by amino acids. Lastly, the changes in Yap signalling caused by acute nutrient deprivation affected the Yap-associated gene Ctgf. Collectively these findings improve our understanding of some of the underlying mechanisms that are involved in Yap signalling in adult cardiomyocytes and may provide insight into potential therapeutic targets which could improve remodelling after MI.

### The Effects of Nutrient Deprivation on Yap Signalling

Yap signalling and methods of regulation have been extensively studied in diverse cell types and disease conditions, but the specific molecular mechanisms underlying the effects of Yap activation after myocardial infarction (MI) are not fully understood. While mechanical stress is considered the primary driver of cardiac Yap signalling [25–27] this study reveals that cardiac metabolic stress can also be a major driver of altered Yap-signalling. The role of Yap-signalling in metabolism has been shown in predominantly cancer metabolism as a regulator of cellular proliferation and substrate utilization [28]. When energy is low, Yap is inhibited by phosphorylation. In times of hypertrophy or proliferation, the absence of essential amino acids could also reasonably act as a checkpoint to Yap-signalling, restricting growth. There is little clarity however on Yap-signaling in the context of non-proliferating cardiomyocytes, or models of the same, and what drives observed changes. Our investigation in cardiomyotubes revealed that acute nutrient deprivation more than any other constituent of ischemia was a salient cause of Yap-phophoregulation and nuclear accumulation. These results are surprising as many other stressors have been implicated as effectors of Yap-signalling in the heart, but this could be due to a lack of cellular specificity amongst a diverse pool of cardiac cells (cardiomyocytes, fibroblasts, vascular cells, et al.). Acute nutrient deprivation did not change total Yap levels, yet it did increase post-translational modification at Yap (p-S397) and Yap (p-S127) in whole cell lysate— consistent with the inactivation and cytoplasmic restriction of Yap signalling, and therefore, not likely to result in increased gene transcription of downstream targets. In determining which essential nutrient was responsible for the acute changes in Yap, initially either glucose and/or fatty acids were postulated as being responsible since they are major energy sources for cardiomyocytes and have previously been linked to phosphorylation of Yap leading to its inactivation [28]. We found that glucose and fatty acid supplementation alone did not reverse the increase in Yap (p-S397) and Yap (p-S127) modifications after nutrient deprivation conditions, thus implicating amino acids as the nutrient responsible. Further analysis of cells with nutrient deprivation revealed a shift in amino acid bioavailability, with non-essential amino acids increased but essential amino acids decreased. This could be explained as essential amino acids by necessity come from extrinsic sources, where non-essential amino acids can be provided by proteolysis, including in excess should the purpose be to extract and recycle essential amino acids. Therefore, in response to acute nutrient deprivation, the cardiomyocytes may quickly consume the essential amino acids to maintain appropriate function and dispense with proteins to build up excess non-essential amino acids thereafter. Of the essential amino acids, threonine was decreased to the greatest extent followed by valine, isoleucine, and leucine—three branched-chain amino acids. Supplementation with threonine and L-isoleucine returned phosphorylated Yap signals to control levels in whole-cell lysate. We then focused on isoleucine as a rate-limiting amino acid, given there exists a feedback loop between threonine and isoleucine by threonine deaminase (the enzyme involved in catalyzing the first reaction which is inhibited by isoleucine allosteric binding to inhibit the reaction at high levels of isoleucine) [21]. As such, increased levels of threonine can be generated by conversion into isoleucine. Additionally, these three branched-chain amino acids were decreased to the same extent, which may be explained by the fact that all three are metabolized by the same enzymes [29]. These findings highlight the critical role of amino acid bioavailability, particularly isoleucine, in modulating Yap signalling in cardiomyocytes under nutrient-deprived conditions.

### Compartmentalization of Yap in Rat Cardiomyotubes After Nutrient Deprivation

Canonically, the translocation of Yap between the nucleus and cytoplasm is a critical mechanism regulating its activity. To investigate the effects of acute nutrient deprivation, we sub-fractionated the nuclear and cytoplasmic compartments of H9c2 cardiomyotubes. Consistent with our whole-cell lysate results, nutrient deprivation increased phosphorylation of Yap at S397 and S127 in both the nucleus and cytoplasm compared to controls. However, this is atypical, as most reports suggest that phosphorylated Yap is cytoplasmically retained, degraded or associated with nuclear export to the cytoplasm [30,31]. It has been suggested that there could be a biphasic pattern of localization for Yap signalling once phosphorylated [32] that is time-dependent [33]. This might account for the increase in total Yap expression in the cytoplasm under nutrient deprivation conditions compared to controls, but our study is time-constrained to just 1hour and we did not account for all post-translational modifications that could occur to Yap and affect binding partners that either promote or restrict gene expression. It may therefore be reasonable to hypothesize that if the time course were expanded, there would be a flux of Yap from the nucleus to the cytoplasm. Interestingly, Yap and Taz do not have a nuclear localization signal which is typically responsible for the accumulation of proteins in the nucleus, therefore the exact mechanism of their nuclear shuttling remains unclear. However, there is evidence that mechanical forces, in addition to biochemical cues, can regulate Yap translocation, particularly in the absence of cell-cell contacts as seen after cardiac injury [34]. Future studies that immunoprecipitate Yap under these conditions could identify key regulatory binding partner proteins necessary for compartmentation or cytoplasmic/nuclear shuttling.

Supplementation with the essential amino acid L-isoleucine maintained Yap, p-Yap(S397), and p-Yap(S127) levels comparable to controls in the whole-cell lysates. Yet, in the subcellular fractions, isoleucine supplementation only returned cytoplasmic p-Yap(S397) to control levels. In the nuclear compartment p-Yap(S397) remained elevated to control. Further, p-Yap(S127) also remained elevated comparatively in both cytoplasm and nuclear fractions even in the presence of isoleucine (Table-1). Typically, phosphorylation at S397 & S127 are associated with inhibition of Yap associated transcription, as it is often associated with the export of Yap to the cytoplasm and subsequently leads to proteasomal degradation [11]. To determine if whole media were able to restore compartmentation to control, cell that were refed normal media for an hour actually accelerated Yap-degradation, and also left S127 Yap elevated as a ratio of total Yap (supplemental figure-1), suggesting that time and substrate are not singular mechanisms operating dependently. Our data are not fully time-resolved and did not account for the canonical or noncanonical sites of regulation. Given that a classically Yap-associated gene, Ctgf, was elevated where S127 was consistently phosphorylated, even in the presence of isoleucine suggests that S127 may be associated with increased gene expression under some circumstances, either by nuclear translocation or retention. Further studies will be required to resolve the nature of Yap shuttling.

**Table 1.**
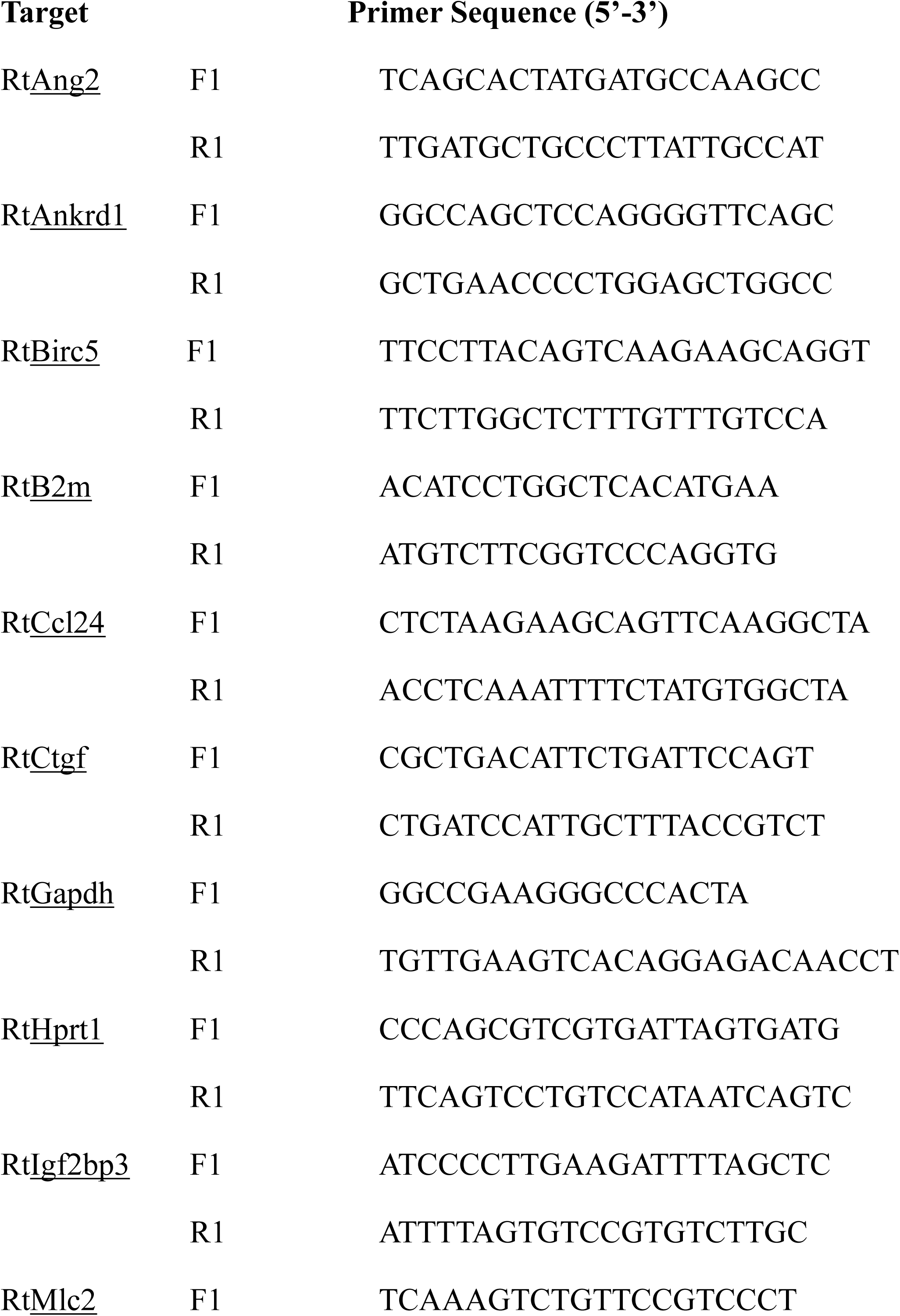
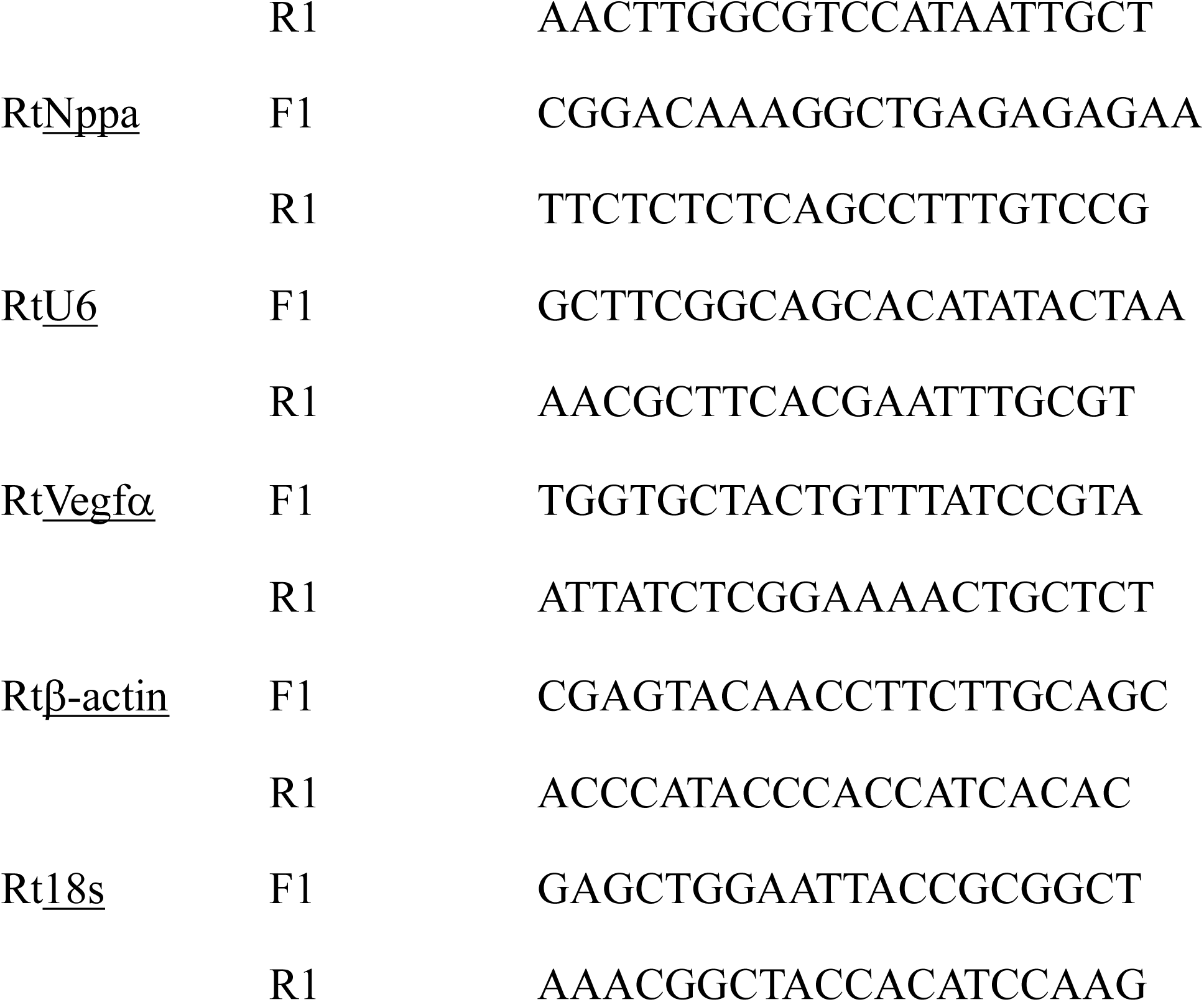
Primer Sequences and Annealing Temperatures of Oligonucleotides used in qPCR.

### Connective Tissue Growth Factor, Yap Signalling and the Heart

The Ctgf gene encodes the Ctgf protein which is also known as cellular communication network factor 2. Regulation of Ctgf can be at the transcriptional, post-transcriptional, and translational levels and is controlled by several pathological and physiological cues. As a secreted protein it is involved in initiating signal transduction by binding to various extracellular constituents and receptors (eg. integrins, heparan sulfate proteoglycans, LRPs, and TrkA). The consequences of Ctgf binding are multifaceted, including effects on the extracellular matrix (ECM), endothelial-mesenchymal transition, macrophage polarization, autophagy, and senescence-associated metabolic disruption [35]. It is responsible for the direct binding of cytokines and mediates ECM-related proteins, in addition to regulating growth factor and cytokine activity by modulating crosstalk between signalling pathways. Ctgf plays a central role in fibrosis by controlling fibroblast proliferation, angiogenesis, matrix production, and ECM deposition. Fibroblasts are fundamental for creating and maintaining the ECM, and their migration, activation, and differentiation into myofibroblasts are major drivers of fibrosis after injury. Ctgf acts as a downstream mediator of transforming growth factor-β (TGF-β), which regulates myofibroblast differentiation and tissue formation. The release of Ctgf from M2 macrophages is also associated with the proliferation and migration of fibroblasts, contributing to ECM formation [36]. In addition to its effects on fibroblasts and macrophages, Ctgf plays a role in endothelial cell function and angiogenesis. Ctgf increases the production of vascular endothelial growth factor (VEGF), which is responsible for angiogenesis, through various pathways (e.g., AKT, ERK and Pi3K) that increase miR-210. Upregulation of miR-210 leads to inhibition of prolyl hydroxylase 2 activity that promotes increased VEGF expression and angiogenesis [37]. The expression of Ctgf is controlled by various transcription factors, including the TEAD transcription factor and its coactivator Yap [35]. Of the Yap-associated genes we tested, only Ctgf expression was markedly changed in response to nutrient deprivation conditions and increased again when L-isoleucine was supplemented back. While there was no change in total Yap expression with isoleucine supplementation, there was a decrease in the inhibitory phosphorylation of Yap at S397 in the cytoplasm compared to nutrient deprivation alone. This suggests that the reduction in Yap phosphorylation may result in the activation of Yap, allowing for the transcription of Ctgf. The activity of Yap/Tead and Ctgf are mutually regulated suggesting that increased activated Yap (unphosphorylated) is correlated with increased Ctgf [126]. The role of Yap in Ctgf-associated angiogenesis was confirmed by re-expression of Yap that rescues angiogenesis in Ctgf mutant mice [38]. This suggests that Ctgf itself may be a feedback regulator of Yap-signalling.

As for fibroblasts, activated Yap increases Ctgf expression to promote fibroblast proliferation [39]. Yap also plays a role in regulating the ECM through Ctgf expression, for example in the heart after MI elevated unphosphorylated Yap coincides with increased ECM production. This was confirmed using verteporfin, a drug which inhibits the binding of Yap to TEAD to arrest gene transcription and decreases ECM production and proliferation of fibroblasts. This therefore resulted in decreased fibrosis and stiffness of the heart after MI compared to controls [40]. Inhibition of Ctgf might improve survival and ejection fraction in addition to improving remodelling by increased cross-sectional area of cardiomyocytes, heart weight and fibrotic genes [41]. Upregulation of Ctgf has also been associated with the hypertrophy of cardiomyocytes [42]. The role of Yap-associated Ctgf is likely cell and time-dependent and may not be readily reversed as we show by simple L-isoleucine supplementation. Its effects are diverse on all cardiac cell types and include both autocrine and paracrine effects likely to influence all major remodelling events after MI but is one of the most rapid responses detected during cardiac stress.

### Clinical Relevance

In a clinical setting, elevated circulating BCAA levels after MI are associated with an increased risk of cardiovascular mortality and acute heart failure. particularly in patients with ST-elevation MI requiring reperfusion [43]. However, this BCAA elevation is not universal, as some studies have found decreased circulating BCAA levels post-MI despite increased cardiac BCAA levels due to impaired BCAA catabolism in mice. *Wang et al.* suggest that permanent MI is correlated with increased cardiac BCAA’s caused by defective catabolism [44]. This impaired BCAA catabolism has been linked to progressive loss of cardiac contractile capacity and early death in animal models [45]. *Gannon et al*. suggest that in conditions where energy is deprived, BCAA might improve the sensitivity to glucose uptake. In conditions where there is excess energy, the catabolism of BCAA is disrupted, which leads to the accumulation of BCAA either in circulation or intracellularly [46]. The regulation of BCAA metabolism appears complex, with evidence that Yap plays an important role, where decreased Yap activity is correlated with lower levels of the amino acid transporter SLC7A5, leading to higher intracellular amino acid levels [47]. This is consistent with our results that show supplementation of L-isoleucine to our nutrient deprivation conditions resulted in decreased Yap (p-S397) in the cytoplasm compared to nutrient deprivation alone and increased Ctgf expression that has been associated with poor remodelling and increased death after MI. Interestingly, Yap activation (unphosphorylated) upregulates SLC7A5 and other nutrient uptake genes, in addition to an upregulation of Ctgf [48]. These findings suggest a potential mechanism between Yap signalling and amino acids may be Yap acting to regulate amino acid metabolism instead of amino acids acting upon Yap signalling, or perhaps it operates in both directions. Given there is a link between elevated BCAA and poor outcomes and activating Yap signalling can decrease amino acid levels by increasing SLC7A5 activity, then increasing Yap signalling after MI may promote better outcomes for patients by improving adaptive remodelling. Therefore, these data highlight a relationship between Yap signalling and amino acids, yet the full mechanism needs to be explored further.

### Limitations and Future Directions

While this paper provides an observational overview of the relationship between Yap signalling and amino acids there are limitations to this work that limit the opportunity to make definitive conclusions or understand the mechanisms fully. Firstly, this study was predominantly done *in vitro* using H9c2 cardiomyocytes. While *in vitro* studies allow for cell specificity and direct modelling of conditions for the identification of molecular mechanisms, it lacks the complexity of the humoral, cellular, and molecular mechanisms interacting amongst various cardiac cell types that are fully matured and functioning to meet physiological demands. Furthermore, our results are from one cell line, comparing our results to another cell line such as AC16’s (human cardiomyocytes) or isolated primary cardiomyocytes could support our results and create a more complete picture of the mechanisms involved and account for different species effects. Furthermore, the short timeline of one hour could be a limitation of our study as it provides only a snapshot in time, it will not resolve what occurs with integrative signalling and temporal progression with MI. While we were able to mimic many drivers of MI in our study, we lacked a model of waste accumulation that occurs after MI, therefore its role in Yap signalling was not identified by this work. Another limitation to our study is that we only measured free amino acid concentrations, however, both circulating and intracellular BCAA play an important role in remodelling after MI. Additionally, our experiments were only carried out testing the supplementation of only three amino acids, and they were all tested separately, combinations of amino acids might influence cell signalling in variable ways we have not determined. Leucine could be of particular interest since isoleucine and leucine have the same chemical formula but slightly different chemical structures. Leucine specifically is implicated in the regulation of other pathways, such as mTOR, and there has been an association between mTOR activity and vascular growth that is YAP/TAZ dependent [48]. Despite these limitations, we have put forward evidence for amino acid-regulated signalling of Yap that can potentially influence remodelling in the context of MI. This is a potentially significant hypothesis-generating study that can lead to further cell-regulated and time-specific in vivo analysis that intersects metabolic changes as both cause and effect in MI outcomes. Moreover, the compartmentation of Yap and its post-translational modifications should be further explored in terms of promoting or restricting gene expression patterns, particularly with unbiased analysis tools.

## Conclusion

Our study provides insights of Yap signalling, showing for the first time that acute nutrient deprivation conditions increase Yap phosphorylation and compartmentation between the nucleaus and cytoplasm. We establish the importance of amino acids as regulators of Yap signalling in a cellular model of cardiomyotubes. Furthermore, our results implicate a relationship between L-isoleucine, Yap and Ctgf gene expression early remodeling where acute nutrient deprivation and amino acid imbalance may occur. Collectively these results should entice further exploration and understanding of the crosstalk between amino acid signaling, including isoleucine, and Yap-mediated remodeling of the heart. Integrating with other signalling cascades could lead to potential therapeutic targets that might improve cardiac remodelling as a novel lead to improve patient outcomes.

## References

1 Thygesen K, Alpert JS, Jaffe AS, et al. Fourth universal definition of myocardial infarction (2018). Russ. J. Cardiol. 2019;24. doi:10.15829/1560-4071-2019-3-107-138

2 Ojha N, Dhamoon AS, Chapagain R. Myocardial Infarction (Nursing). 2021.

3 Burke AP, Virmani R. Pathophysiology of acute myocardial infarction. Med Clin North Am 2007;91:553–72; ix. doi:10.1016/j.mcna.2007.03.005

4 Chen R, Suchard MA, Krumholz HM, et al. Comparative First-Line Effectiveness and Safety of ACE (Angiotensin-Converting Enzyme) Inhibitors and Angiotensin Receptor Blockers: A Multinational Cohort Study. Hypertens (Dallas, Tex 1979) 2021;78:591–603. doi:10.1161/HYPERTENSIONAHA.120.16667

5 Del Re DP, Amgalan D, Linkermann A, et al. Fundamental mechanisms of regulated cell death and implications for heart disease. Physiol Rev 2019;99. doi:10.1152/physrev.00022.2018

6 Nakamura M, Sadoshima J. Mechanisms of physiological and pathological cardiac hypertrophy. Nat. Rev. Cardiol. 2018;15. doi:10.1038/s41569-018-0007-y

7 Nauta JF, Hummel YM, Tromp J, et al. Concentric vs. eccentric remodelling in heart failure with reduced ejection fraction: clinical characteristics, pathophysiology and response to treatment. Eur J Heart Fail 2020;22. doi:10.1002/ejhf.1632

8 Kim W, Jho EH. The history and regulatory mechanism of the Hippo pathway. BMB Rep. 2018;51. doi:10.5483/BMBRep.2018.51.3.022

9 Pocaterra A, Romani P, Dupont S. YAP/TAZ functions and their regulation at a glance. J Cell Sci 2020;133. doi:10.1242/jcs.230425

10 Kim MK, Jang JW, Bae SC. DNA binding partners of YAP/TAZ. BMB Rep. 2018;51. doi:10.5483/BMBRep.2018.51.3.015

11 Wang P, Gong Y, Guo T, et al. Activation of Aurora A kinase increases YAP stability via blockage of autophagy. Cell Death Dis 2019;10. doi:10.1038/s41419-019-1664-4

12 Moon S, Kim W, Kim S, et al. Phosphorylation by NLK inhibits YAP -14-3-3- interactions and induces its nuclear localization. EMBO Rep 2017;18. doi:10.15252/embr.201642683

13 Deng F, Wu Z, Zou F, et al. The Hippo-YAP/TAZ Signaling Pathway in Intestinal Self-Renewal and Regeneration After Injury. Front cell Dev Biol 2022;10:894737. doi:10.3389/fcell.2022.894737

14 Hora S, Wuestefeld T. Liver Injury and Regeneration: Current Understanding, New Approaches, and Future Perspectives. Cells 2023;12. doi:10.3390/cells12172129

15 Wang J, Liu S, Heallen T, et al. The Hippo pathway in the heart: pivotal roles in development, disease, and regeneration. Nat. Rev. Cardiol. 2018;15:672–84. doi:10.1038/s41569-018-0063-3

16 Xiao Y, Leach J, Wang J, et al. Hippo/Yap Signaling in Cardiac Development and Regeneration. Curr. Treat. Options Cardiovasc. Med. 2016;18. doi:10.1007/s11936-016-0461-y

17 Xin M, Kim Y, Sutherland LB, et al. Hippo pathway effector Yap promotes cardiac regeneration. Proc Natl Acad Sci U S A 2013;110:13839–44. doi:10.1073/pnas.1313192110

18 Gong R, Jiang Z, Zagidullin N, et al. Regulation of cardiomyocyte fate plasticity: a key strategy for cardiac regeneration. Signal Transduct Target Ther 2021;6:31. doi:10.1038/s41392-020-00413-2

19 Brazma A, Hingamp P, Quackenbush J, et al. Minimum information about a microarray experiment (MIAME) - Toward standards for microarray data. Nat. Genet. 2001;29. doi:10.1038/ng1201-365

20 Tulacz D, Mackiewicz U, Maczewski M, et al. Transcriptional profiling of left ventricle and peripheral blood mononuclear cells in a rat model of postinfarction heart failure. BMC Med Genomics 2013;6. doi:10.1186/1755-8794-6-49

21 Isogai S, Nishimura A, Kotaka A, et al. High-Level Production of Isoleucine and Fusel Alcohol by Expression of the Feedback Inhibition-Insensitive Threonine Deaminase in Saccharomyces cerevisiae. Appl Environ Microbiol 2022;88. doi:10.1128/aem.02130-21

22 Lin Z, Von Gise A, Zhou P, et al. Cardiac-specific YAP activation improves cardiac function and survival in an experimental murine MI model. Circ Res 2014;115. doi:10.1161/CIRCRESAHA.115.303632

23 Heallen T, Morikawa Y, Leach J, et al. Hippo signaling impedes adult heart regeneration. Dev 2013;140. doi:10.1242/dev.102798

24 Correction (JACC: Basic to Translational Science (2020) 5(9) (931–945), (S2452302X20303417), (10.1016/j.jacbts.2020.07.009)). JACC Basic to Transl. Sci. 2021;6. doi:10.1016/j.jacbts.2021.06.007

25 Wang J, Liu S, Heallen T, et al. The Hippo pathway in the heart: pivotal roles in development, disease, and regeneration. Nat Rev Cardiol 2018;15:672–84. doi:10.1038/s41569-018-0063-3

26 Wang M, Lin BY, Sun S, et al. Shear and hydrostatic stress regulate fetal heart valve remodeling through YAP-mediated mechanotransduction. Elife 2023;12. doi:10.7554/eLife.83209

27 Wong DCP, Xiao J, Chew TW, et al. BNIP-2 Activation of Cellular Contractility Inactivates YAP for H9c2 Cardiomyoblast Differentiation. *Adv Sci (Weinheim, Baden-Wurttemberg*, Ger 2022;9:e2202834. doi:10.1002/advs.202202834

28 Koo JH, Guan KL. Interplay between YAP/TAZ and Metabolism. Cell Metab. 2018;28. doi:10.1016/j.cmet.2018.07.010

29 Dimou A, Tsimihodimos V, Bairaktari E. The Critical Role of the Branched Chain Amino Acids (BCAAs) Catabolism-Regulating Enzymes, Branched-Chain Aminotransferase (BCAT) and Branched-Chain α-Keto Acid Dehydrogenase (BCKD), in Human Pathophysiology. Int. J. Mol. Sci. 2022;23. doi:10.3390/ijms23074022

30 Wada KI, Itoga K, Okano T, et al. Hippo pathway regulation by cell morphology and stress fibers. Development 2011;138. doi:10.1242/dev.070987

31 Dupont S, Morsut L, Aragona M, et al. Role of YAP/TAZ in mechanotransduction. Nature 2011;474. doi:10.1038/nature10137

32 Wang J, Sinnett-Smith J, Stevens J V., et al. Biphasic regulation of Yes-associated Protein (YAP) cellular localization, phosphorylation, and activity by G proteincoupled receptor agonists in intestinal epithelial cells: A novel role for Protein Kinase D (PKD). J Biol Chem 2016;291. doi:10.1074/jbc.M115.711275

33 Shreberk-Shaked M, Oren M. New insights into YAP/TAZ nucleo-cytoplasmic shuttling: new cancer therapeutic opportunities? Mol. Oncol. 2019;13. doi:10.1002/1878-0261.12498

34 Das A, Fischer RS, Pan D WC. YAP Nuclear Localization in the Absence of Cell-Cell Contact Is Mediated by a Filamentous Actin-dependent, Myosin II-and Phospho-YAP-independent Pathway during Extracellular Matrix Mechanosensing. J Biol Chem 2016;291.

35 Effendi WI, Nagano T. Connective Tissue Growth Factor in Idiopathic Pulmonary Fibrosis: Breaking the Bridge. Int. J. Mol. Sci. 2022;23. doi:10.3390/ijms23116064

36 Zhang SM, Wei CY, Wang Q, et al. M2-polarized macrophages mediate wound healing by regulating connective tissue growth factor via AKT, ERK1/2, and STAT3 signaling pathways. Mol Biol Rep 2021;48. doi:10.1007/s11033-021-06646-w

37 Liu SC, Chuang SM, Hsu CJ, et al. CTGF increases vascular endothelial growth factor-dependent angiogenesis in human synovial fibroblasts by increasing miR-210 expression. Cell Death Dis 2014;5. doi:10.1038/cddis.2014.453

38 Moon S, Lee S, Caesar JA, et al. A CTGF-YAP Regulatory Pathway Is Essential for Angiogenesis and Barriergenesis in the Retina. iScience 2020;23. doi:10.1016/j.isci.2020.101184

39 Sharifi-sanjani M, Berman M, Goncharov D, et al. Yes-associated protein (Yap) is up-regulated in heart failure and promotes cardiac fibroblast proliferation. Int J Mol Sci 2021;22. doi:10.3390/ijms22116164

40 Garoffolo G, Casaburo M, Amadeo F, et al. Reduction of Cardiac Fibrosis by Interference with YAP-Dependent Transactivation. Circ Res 2022;131. doi:10.1161/CIRCRESAHA.121.319373

41 Vainio LE, Szabó Z, Lin R, et al. Connective Tissue Growth Factor Inhibition Enhances Cardiac Repair and Limits Fibrosis After Myocardial Infarction. JACC Basic to Transl Sci 2019;4. doi:10.1016/j.jacbts.2018.10.007

42 Matsui Y, Sadoshima J. Rapid upregulation of CTGF in cardiac myocytes by hypertrophic stimuli: Implication for cardiac fibrosis and hypertrophy. J. Mol. Cell. Cardiol. 2004;37. doi:10.1016/j.yjmcc.2004.05.012

43 Du X, You H, Li Y, Wang Y, Hui P, Qiao B, Lu J, Zhang W, Zhou S, Zheng Y DJ. Relationships between circulating branched chain amino acid concentrations and risk of adverse cardiovascular events in patients with STEMI treated with PC. Sci Rep 2018;8.

44 Wang W, Zhang F, Xia Y, et al. Defective branched chain amino acid catabolism contributes to cardiac dysfunction and remodeling following myocardial infarction. Am J Physiol - Hear Circ Physiol 2016;311. doi:10.1152/ajpheart.00114.2016

45 Wendel U. Metabolism of branched-chain amino acids in maple syrup urine disease. Eur J Pediatr Suppl 1997;156. doi:10.1007/pl00014274

46 Gannon NP, Schnuck JK, Vaughan RA. BCAA Metabolism and Insulin Sensitivity – Dysregulated by Metabolic Status? Mol. Nutr. Food Res. 2018;62. doi:10.1002/mnfr.201700756

47 Najumudeen AK, Ceteci F, Fey SK, et al. The amino acid transporter SLC7A5 is required for efficient growth of KRAS-mutant colorectal cancer. Nat Genet 2021;53. doi:10.1038/s41588-020-00753-3

48 Ong YT, Andrade J, Armbruster M, et al. A YAP/TAZ-TEAD signalling module links endothelial nutrient acquisition to angiogenic growth. Nat Metab 2022;4. doi:10.1038/s42255-022-00584-y

