## Supplemental Figures for "Yap is a Nutrient Sensor Sensitive to the Amino Acid L-Isoleucine and Regulates Expression of Ctgf in Cardiomyocytes"

**Supplemental Figure 1**

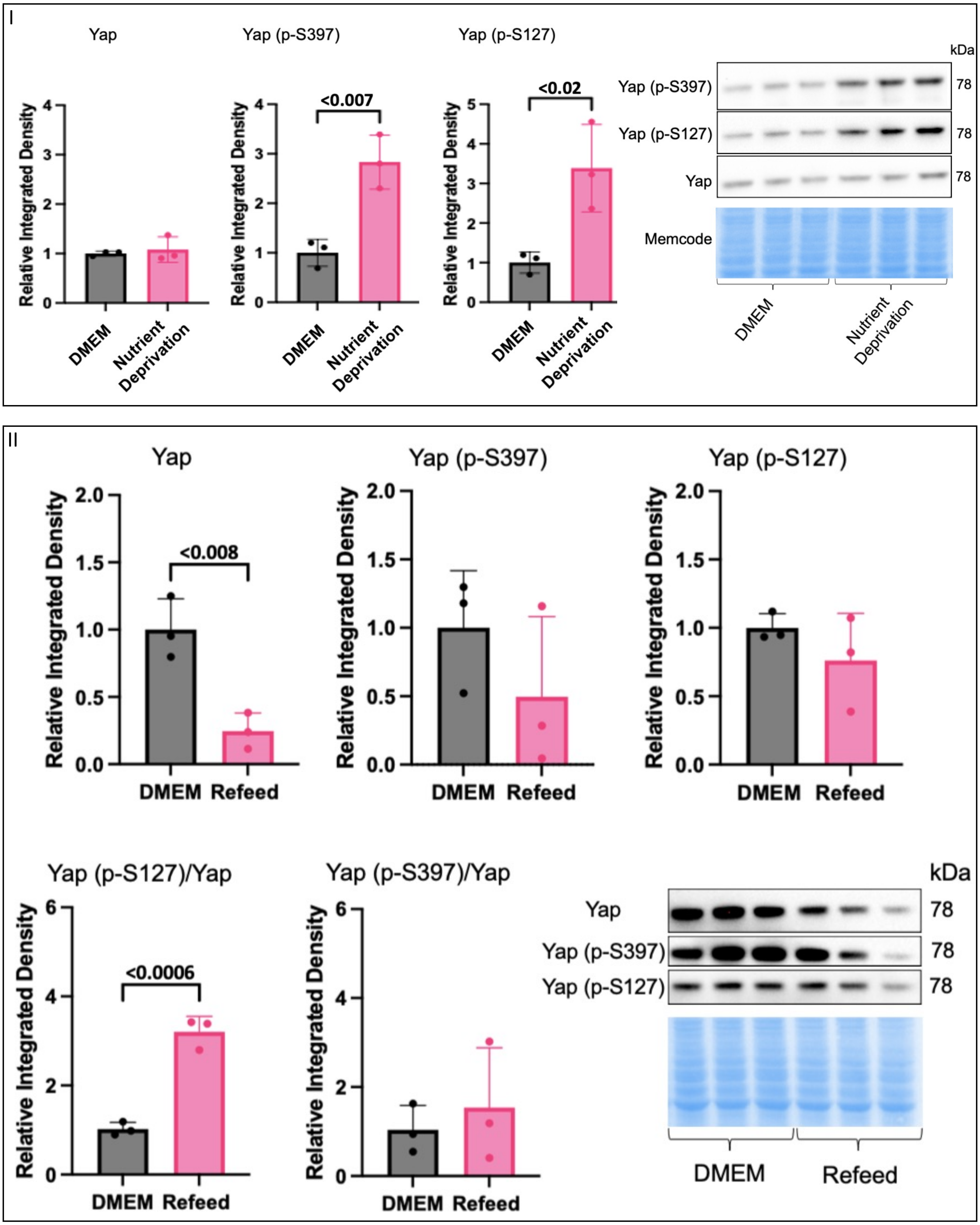


**Supplemental Figure 1. Effects of Nutrient Deprivation on Yap Signaling.** H9c2 cells were differentiated for 6 days in the absence of FBS, on the day of the experiment controls received a media change and our treatment groups received nutrient deprivation solution for 1 hour before harvest. **(I)** Yap expression was unchanged compared to controls. Yap (p-S397) was significantly increased in nutrient deprivation conditions compared to controls. Yap (p-S127) was also significantly increased under nutrient deprivation conditions compared to controls. Representative Yap, Yap (p-S397), and Yap (p-S127) blots with respective Pierce® Memcode demonstrating uniform protein loading. II) On the day of the experiment controls received a media change and our treatment groups received nutrient deprivation solution for one hour, followed by re-addition of DMEM control media for one hour. Refeeding with DMEM decreased total Yap expression after one hour but had no effect on Yap (p-S397) and Yap (p-S127) alone. Ratioed Yap (p-S127) to total Yap expression was increased in the refeed group compared to DMEM controls. Ratioed Yap (p-S397) to total Yap expression had no significant changes compared to controls**.** Representative blots for Yap, Yap (p-S397) and Yap (p-S127) and Pierce® Memcode demonstrating uniform protein loading

**Supplemental Figure 2**

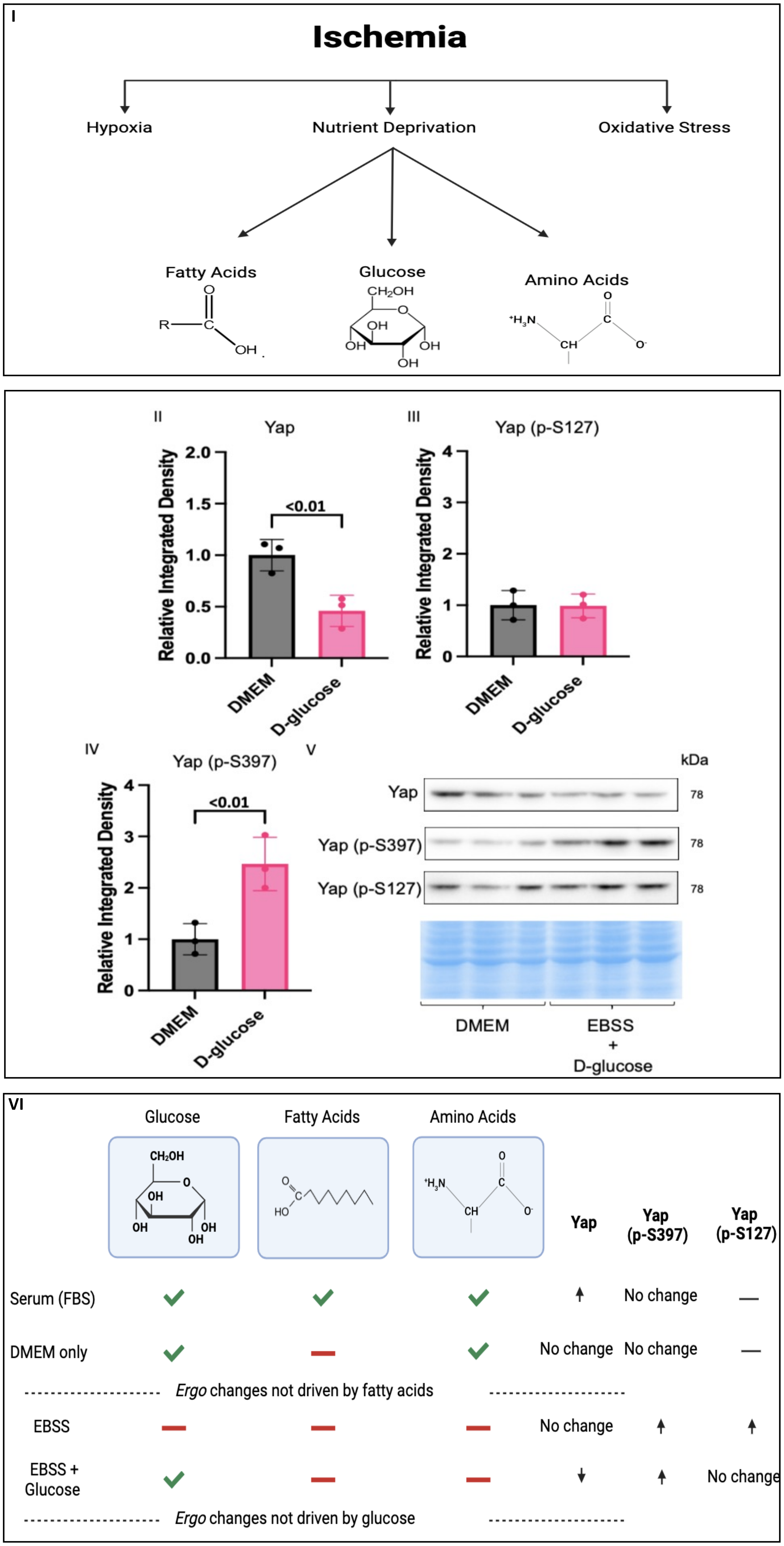

**Supplemental Figure 2. Ischemic components with Glucose Supplementation. I)** Ischemia is complex and has many components, three major components are hypoxia, oxidative stress, and nutrient deprivation. We have previously ruled out hypoxia and oxidative stress for changes in phosphorylation patterns for Yap, therefore, nutrient deprivation was left to test. Nutrient deprivation constitutes of deprivation of three essential nutrients, fatty acids, glucose, and amino acids and each component will be tested separately to determine their effects on Yap signaling. H9c2 cells were differentiated for 6 days in the absence of FBS, on the day of the experiment controls received a media change and our treatment groups received metabolic shock solution supplement with 4500g/L D-glucose for 1 hour. **(II)** D-glucose supplementation decreased Yap expression compared to DMEM. (I**II)** There were no changes in Yap (p-S127) compared to DMEM. **(IV)** Yap (p-S397) was increased in D-glucose groups compared to DMEM. **(V)** Respective Yap, Yap (p-S127) and Yap (p-S397) blots with Pierce® Memcode as protein loading control. VI) Schematic depicting the changes or lack thereof in Yap, Yap (p-S397) and Yap (p-S127) caused by FBS (fatty acids) which allowed us to determine that fatty acids did not cause changes in Yap phosphorylation. Both Yap S397 and S127 were increased in nutrient deprivation (EBSS) while there were no changes in total Yap expression. Supplementation with D-glucose decreased total Yap expression compared to DMEM controls, while Yap (p-S397) was increased with no change in Yap (p-S127). Schematic depicting the changes or lack thereof in Yap, Yap (p-S397) and Yap (p-S127) caused by FBS (fatty acids) which allowed us to determine that fatty acids did not cause changes in Yap phosphorylation. Both Yap S397 and S127 were increased in metabolic shock (EBSS) while there were no changes in total Yap expression. Supplementation with D-glucose decreased total Yap expression compared to DMEM controls, while Yap (p-S397) was increased with no change in Yap (p-S127).

**Supplemental Figure 3**

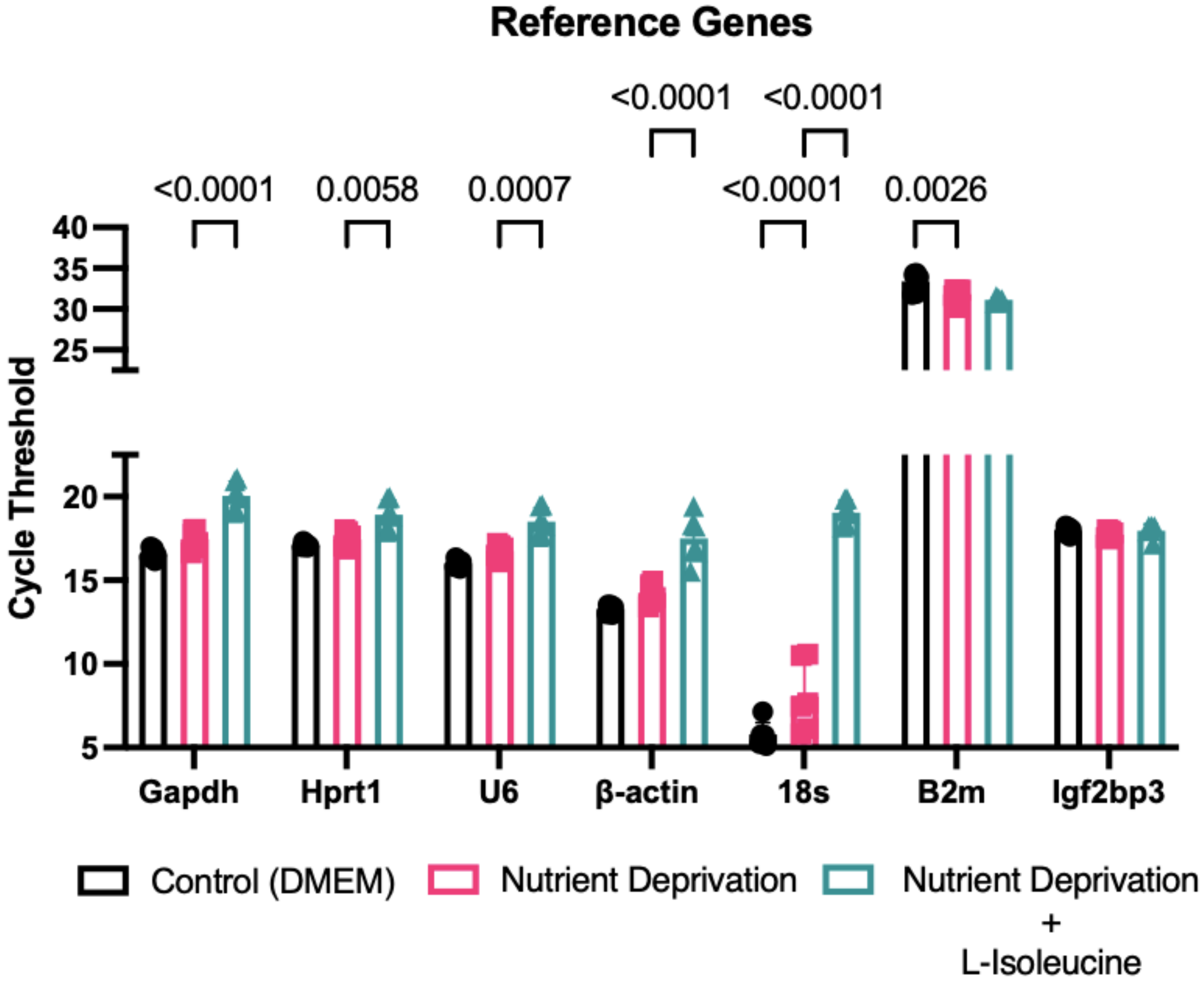


**Supplemental Figure 3. Determination of Reference Genes for qPCR Analysis.** H9c2 cells were differentiated for 6 days in the absence of FBS, on the day of the experiment controls received a media change and our treatment groups received nutrient deprivation solution or nutrient deprivation with L-isoleucine for 1 hour. RNA was extracted and converted to cDNA which was used for qPCR. Six common housekeeping (reference genes) used for H9c2 cells were tested, however all were significantly different between one or more groups. Igf2bp3 was the only gene tested that did not change between all three groups.
